# Proteomic and Metabolomic Profiling of Archaeal Extracellular Vesicles from the Human Gut

**DOI:** 10.1101/2024.06.22.600174

**Authors:** Viktoria Weinberger, Barbara Darnhofer, Polona Mertelj, Regis Stentz, Himadri B Thapa, Emily Jones, Gerlinde Grabmann, Rokhsareh Mohammadzadeh, Tejus Shinde, Rokas Juodeikis, Dominique Pernitsch, Kerstin Hingerl, Tamara Zurabishvili, Christina Kumpitsch, Torben Kuehnast, Dagmar Kolb, Kathryn Gotts, Thomas Weichhart, Thomas Köcher, Harald Köfeler, Simon R. Carding, Stefan Schild, Christine Moissl-Eichinger

**Affiliations:** Diagnostic and Research Institute of Hygiene, Microbiology and Environmental Medicine, Medical University of Graz, Austria; Core Facility Mass Spectrometry, Medical University of Graz, Graz, Austria; Food, Microbiome and Health Institute Research Programme, Quadram Institute Bioscience, Norwich, United Kingdom; Institute of Molecular Biosciences, University of Graz, Graz, Austria; Vienna BioCenter Core Facilities GmbH Metabolomics, Vienna, Austria; Core Facility Ultrastructure Analysis, Medical University of Graz, Graz, Austria; Core Science Resources, Quadram Institute Bioscience, Norwich, United Kingdom; Center for Pathobiochemistry and Genetics, Medical University of Vienna, Vienna, Austria; Norwich Medical School, University East Anglia, Norwich, United Kingdom; Field of Excellence Biohealth – University of Graz, Graz, Austria; BioTechMed Graz, Austria

## Abstract

One potential mechanism for microbiome-host, and microbiome constituents’ interaction and communication involves extracellular vesicles (EVs). Here, for the first time, we report the capability of two M. smithii strains (ALI and GRAZ-2), Candidatus M. intestini, and Methanosphaera stadtmanae, as underrepresented components of the gut microbiome, to produce EVs. Interesting, size, morphology, and composition of AEVs were comparable to bacterial EVs, as indicated by ultrastructure, composition, proteomic and metabolomic analyses; however, EVs were substantially less prevalent in the studied Archaea. When looking at the proteomics more precisely, although AEVs from M. smithii ALI and M. intestini were found to be carrying unique proteins (n=135 and n=30, respectively), the shared proteins in AEVs within this genus (n=229), were mostly adhesins(/like) proteins, or proteins with IG-like domains. One remarkable observation was the uptake of AEVs obtained from Methanosphaera stadtmanae and the studied Methanobrevibacter species by human monocytes and the subsequent IL-8 secretion.

## Introduction

All organisms have evolved various signaling mechanisms to convey crucial biological information across cells, tissues, and organs^1–3^. Among these mechanisms are extracellular vesicles (EVs), which are small membrane-bound spherical particles produced and released by cells of all three domains of life^1^.

In the gastrointestinal tract (GIT), extracellular vesicles produced by commensal bacteria (bacterial extracellular vesicles, BEVs) mediate intra and inter-kingdom interactions, maintaining the microbiome ecosystem and promoting interactions with the host ^2^.

BEVs have garnered considerable attention in recent years due to their diverse roles in intercellular communication, pathogenesis, stress tolerance, immune stimulation, and host-microbe interactions^3–7^. These small, membrane-bound structures serve as vehicles for the transport of biomolecules such as proteins, nucleic acids, metabolites and lipids between bacterial cells, as well as between bacteria and their host environments ^4,8–12^. Understanding the mechanisms underlying BEV biogenesis, cargo loading, and their impact on microbial communities and host physiology is critical in microbiology and biomedical research ^4^.

BEVs are divided into different categories based on either their producing bacteria (BEVs from Gram-negative and Gram-positive bacteria) or their origin and the pathway by which they are formed (outer membrane vesicles, outer-inner membrane vesicles, explosive membrane vesicles or cytoplasmic membrane vesicles)^13,14^.

Outer membrane vesicles (OMVs) are considered as the archetypal bacterial membrane vesicles. These OMVs usually arise from a protrusion of the outer membrane and their envelope, therefore resembling the envelope of the donor cell. They usually contain surface-associated factors, outer membrane proteins, and periplasmic content^13,14^. Explosive membrane vesicles on the other hand diversify BEV composition, explaining the presence of nucleic acids and cytosolic content in vesicle samples from Gram-negative bacteria.

In the course of the last decade, it has become evident that BEVs of GIT-colonizing bacteria potentially influence essential functions of the intestine, and of systemic organs after their migration to the bloodstream, thereby contributing to host health^15^. For instance, BEVs contribute to host digestion by distributing hydrolase activities across the lumen, and can potentially influence the central nervous system following migration through the gut-brain axis^16^. Additionally, BEVs can act as efficient delivery vehicles of bioactive compounds, such as toxins or modulators of host cell physiology^13,14^. BEVs are recognized and efficiently internalized by various host cells resulting in intestinal barrier changes, immunomodulation and (patho-)physiological changes^13,14^. BEVs can also act on the surrounding microbiota, promoting bacterial colonization and growth as well as protecting bacteria from antibiotics and host defense peptides ^11,17,18^.

Triggers for vesicle formation are manifold, including factors such as media composition, growth phase, temperature, iron and oxygen availability, as well as exposure to antibiotics and stress^13,14^. As a consequence of the diverse triggers and various origins, the vesicle preparations likely reflect a mixture of different BEV types, which could explain variable BEV functions and effects, and experiments are sometimes non-conclusive^14^.

Representatives of all three domains of eukaryotes, bacteria, and archaea, are capable of forming extracellular vesicles^19^. The reports on archaeal vesicles are overall fairly rare and are restricted to extremophilic archaea, namely Thermococcales and Sulfolobales. It appears that in *Sulfolobus*, for example, vesicle formation is evolutionarily related to eukaryotic ESCRT complex proteins used for the building of endosomes; however, other archaea, such as *Thermococcus* form vesicles but do lack the ESCRT complex, indicating a higher variety in vesicle formation mechanisms^19^. In general, a defensive function through these vesicles was proposed, but research is still ongoing^20^. However, archaea not only thrive in environmental ecosystems, but are also considered as reliable and prevalent constituents of the human GIT microbiome. With 1.2% relative abundance on average, *Methanobrevibacter* and *Methanosphaera* species are highly prevalent across individuals (>90%)^21,22^. Trough maintaining numerous syntrophic relationships with intestinal bacteria, these archaea have the capacity to orchestrate the entire microbiome, leading to an optimized fibre degradation^23^. They also influence the host with respect to the provision of short chain fatty acids or mediate the reduction of gut motility, leading to constipation^18^. However, the mechanisms by which they interact with other microorganisms and their mode of signaling have remained unknown. In this manuscript, we focus on the recent discovery of archaeal extracellular vesicles (AEVs) produced by human archaeal representatives and present novel findings on their ultrastructure, proteome, and metabolome, as well as their interaction with human cell lines. We will discuss the implications of this discovery for our understanding of microbiome-host interactions and outline future directions for research. By integrating insights from bacterial and archaeal EV biology, we strive to unravel the complexities of microbial communication networks within the human body and their implications for health and disease.

## Material and Methods

### Source of microorganisms

The human gut derived strains *Methanobrevibacter smithii* ALI (DSM 2375), and *Methanosphaera stadtmanae* (DSM 3091, type strain) were obtained from the German Collection of Microorganisms and Cell Cultures (DSMZ) GmbH, Braunschweig, Germany. *Candidatus* M. intestini WWM1085 (DSM 116060) was obtained from the Department of Microbiology, University of Illinois, USA, where it was isolated from a stool sample of a healthy woman^24^. In the following, we will use the abbreviation “M. intestini” instead of *Candidatus* M. intestini.

*M. smithii* GRAZ-2 (DSM 116045) was isolated in 2018 at the Medical University of Graz, Graz, Austria, from a stool sample of a healthy woman^24^. Instead of opting for the *Methanobrevibacter smithii* type strain (PS, DSM 861), our choice was *M. smithii* ALI, as it sourced from a human fecal sample and not from sewage water. Enterotoxigenic *Escherichia coli* (ETEC) H10407 and *Bacteroides fragilis* ATCC® 25285 have been reported previously^25^.

### Growth media and cultivation

For the cultivation of all methanogens standard methanogenium medium (MS) with some modifications as previously described ^24^. For vesicle production, aliquots of 250 ml media in 1000 ml infusion bottles were sealed, pressurized with H_2_/CO_2_ (4:1) and autoclaved. Before inoculation and incubation at 37°C, sodium acetate (0.001g/ml, anoxic, sterile) and yeast extract (0.001g/ml, anoxic, sterile, YE) were added to the media. Vesicles of ETEC and *B. fragilis* were retrieved from stocks prepared earlier ^25^.

### Electron microscopy

Electron microscopy (EM) was undertaken at the Core Facility Ultrastructure Analysis, Medical University of Graz, Graz, Austria and at the Core Science Resources Quadram Institute Bioscience, Norwich, United Kingdom. For ultrastructural analyses of cells, isolates were cultivated in 20 ml aliquots in 100 ml serum bottles for 7 days under anaerobic conditions at 37°C in an incubation shaker (shaking speed: 80 rpm). Followed by the centrifugation of 2 ml of medium containing each strain at 4000 g, 4°C, for 10 min. Cell pellets were then directly handed over to the Core Facility Ultrastructures, Medical University Graz, Graz, Austria for further preparation. AEVs (1x10^11^/ml) were directly handed over to the Core Science Resources Quadram Institute Bioscience, Norwich, United Kingdom.

### Transmission electron microscopy: thin sections and tomography

Cells were fixed in 2.5% (w/v) glutaraldehyde and 2% (w/v) paraformaldehyde in 0.1 M cacodylate buffer, pH 7.4, for 1 h, postfixed in 1% (w/v) osmium tetroxide for 2 h at room temperature, dehydrated in graded series of ethanol and embedded in TAAB (Agar Scientific, Essex, GB) epoxy resin. Ultrathin sections (70 nm thick) were cut with a UC 7 Ultramicrotome (Leica Microsystems, Vienna, Austria) and stained with lead citrate for 5 min and with platinum blue for 15 min. Images were taken using a Tecnai G2 20 transmission electron microscope (Thermo Fisher) with a Gatan ultrascan 1000 charge coupled device (CCD) camera (temperature −20 °C; acquisition software Digital Micrograph; Gatan, Munich, Germany). The acceleration voltage was 120 kV. The tilt series was reconstructed using FLARA, a joint alignment and reconstruction algorithm for electron tomography. This iterative algorithm allows for acquisitions without fiducial gold markers, since an effective shift computation can be obtained by using a global alignment technique based on a linearized approximation of the disruptive shifts in each iteration^26^. For negative staining cell suspensions were placed on glow discharged carbon coated copper grids for 1 min. The solution was removed after incubation by filter paper stripes. A drop of 1% aqueous uranyl acetate solution was placed afterwards for 1 min, dried with filter paper and later on air dried at room temperature. Specimens were examined with an FEI Tecnai G 2 (FEI, Eindhoven, Netherlands) equipped with a Gatan ultrascan 1000 charge coupled device (CCD) camera (-20°C, acquisition software Digital Micrograph, Gatan, Munich, Germany).

AEV suspensions were visualized using negative staining with TEM. Briefly, 4 μL AEV suspension was adsorbed to plasma-pretreated carbon-coated copper EM grids (EM Solutions) for 1 min before wicking off with filter paper and negatively staining with 1% Uranyl Acetate solution (BDH 10288) for 1 min. Grids were air-dried before analysis using a FEI Talos F200C electron microscope at 36,000×-92,×000 magnification with a Gatan OneView digital camera.

### Scanning electron microscopy

For scanning electron microscopy, cells were affixed to coverslips and treated with a fixing solution consisting of 2% paraformaldehyde and 2.5% glutaraldehyde in 0.1 M phosphate buffered saline (pH 7.4). Subsequently, a graded ethanol series was used for dehydration. Post-fixation involved 1% Osmium tetroxide for 1 hour at room temperature, followed by additional dehydration in an ethanol series (ranging from 30% to 100% EtOH). Hexamethyldisilazane (HMDS) was applied, and coverslips were positioned on stubs using conductive double-coated carbon tape. Imaging was performed with a Sigma 500VP FE-SEM equipped with a SEM Detector (Zeiss Oberkochen) operating at an acceleration voltage of 5 kV.

### AEV Isolation

To obtain a sufficient amount of biomass for the isolation of AEVs, 250 ml of MS medium was aliquoted into 1000 ml infusion bottles (VWR) and further handled the same way as described above. These cultures were then cultivated for 10 days under anaerobic conditions at 37°C in an incubation shaker (shaking speed: 80 rpm). When the pressure of cultivation bottles dropped due to growth, they were re-gassed with H2/CO2. Growth was surveyed by optical density photometry at 600 nm. On day ten, the cell suspensions were centrifuged at 14,000 x g, 4°C, 20 min (Thermo Scientific™ Sorvall™ LYNX™ 6000). To remove cell debris and remaining cells, the supernatant was filtered with 0.22 µm PES bottle-top filters (Fisherbrand™ Disposable PES Bottle Top Filters). If not immediately processed, the supernatant containing the vesicles was stored at 4°C.

Isolation of vesicles was done according to Stentz et al.^27^ (Workflow see Supplementary Fig S1). In brief, a filtration cassette (Vivaflow 50R, 100,000 MWCO, Hydrostat, model VF05H4, Sartorius or Vivaflow 200 100,00 MWCO, PES, model VF20P4) was used to concentrate 1 L of sample down to approx. 5 ml. Then, 500 ml PBS buffer (pH 7.4) was added for washing purposes, and the liquid was concentrated to 1-4 ml. The sample was then centrifuged for 20 min at 10,000 g, 4°C to remove protein and lipid aggregates. Next, the sample was transferred to Pierce™ Protein Concentrators (PES, 100,000 MWCO, Thermo Scientific) and centrifuged at 3,000 g until the samples were concentrated down to 1 ml. Residual contaminants and proteins were further eliminated through size exclusion chromatography (SEC) using an IZON qEV1 column (pore size 35 mm) according to the manufacturer’s instructions. The vesicles were eluted in the 2.8 ml fraction containing the purified extracellular vesicles underwent a final filter sterilization using a 0.22 µm syringe filter (ROTILABO® PES, 0,22 µm), and were subsequently stored at 4°C until further use.

To ensure that the final AEV suspension does not contain any yeast vesicles or other residues, the YE was sterile-filtered previous to medium preparation.

For the metabolomics analyses, 1 L of blank MS medium underwent the same procedure to serve as a control.

### BEV Isolation

BEVs for the HT-29 experiment were isolated as described previously with minor modifications^25,28^. Briefly, overnight cultures were either grown with aeration (180 rpm, Infor shaker) in case of ETEC or anaerobically (GasPak™ EZ Systems, BD) in case of *B. fragilis* to ensure sufficient growth. The respective cultures were diluted (1:100) in BHI medium and grown at 37°C either with aeration for 8 h or overnight anaerobically (GasPak™ EZ Systems, BD). The cells were then removed from the supernatant by centrifugation (9,000 x g, 15 min) and subsequent sterile filtration (0.22 µm). The BEVs present in the supernatant were pelleted through subsequent ultracentrifugation (150,000 x g, 4°C, 4 h), resuspended in appropriate volumes of PBS to generate a BEV suspension 1000-fold more concentrated than in the original culture supernatant. Quantification and size distribution of BEVs were investigated by nanoparticle tracking analysis (NTA) using a Nanosight NS300 (see below).

### AEV characterization

#### Nanoparticle tracking analysis (NTA)

Quantification and size distribution of AEVs were investigated by nanoparticle tracking analysis (NTA) using ZetaView and Nanosight NS300. ZetaView was used by following established protocols^27,29^. In brief, particles were quantified using the ZetaView instrument (Particle Metrix, Germany) with ZetaView (version 8.05.12 SP1) software running a 2 cycle 11 position high frame rate analysis at 25°C. Samples were diluted with ultrapure water allowing the optimal detection range. Camera control settings: 80 Sensitivity; 30 Frame Rate; 100 Shutter. Post-acquisition parameters: 20 Min Brightness; 2000 Max Area; 5 Min Area; 30 Trace Length; 5 nm/Class; 64 Classes/Decade.

For NanoSight NS300 (Malvern Instruments, UK) samples were diluted in 1x PBS according to the manufacturer’s guidelines (final concentration between 10^7^ - 10^9^ particles per ml), and a 405 nm laser was used. Between samples, the instrument was flushed with 10% Ethanol and Aqua.dest. Reads of 1-minute duration were performed in five replicates for each sample with the following capture settings: cell temperature: 25°C, syringe load/flow rate: 30, camera: sCMOS. For capture settings, camera level was adjusted so that all particles were distinctly visible (Camera level 12 - 15). The ideal detection threshold was set including as many particles as possible and debris (blue cross count) with a maximum of five (detection threshold 5). Data output was acquired using NanoSight NTA software version 3.3 (Malvern Instruments). For each sample, the mean particle number in the Experiment Summary output was adjusted by the dilution factor.

#### Protein, DNA, and RNA content

As previously described^30–34^, quantification of vesicle content, including protein, DNA, and RNA, was conducted using the Qubit^®^ Protein Assay, Qubit^®^ dsDNA high sensitivity assay, and RNA high sensitivity assay kits, respectively (Thermo Fisher Scientific). Protein, DNA, and RNA measurements were performed using a Qubit^®^ 4 or Qubit^®^ 3 Fluorometer. Instructions of the manufacturer were followed.

#### Lipid content

The quantification of lipid content in AEVs was conducted using the FM4-64 lipophilic fluorescent dye and a linoleic acid standard, a method previously employed for bacterial extracellular vesicle (BEV) lipid quantification^35^. The modified procedure for quantifying vesicles released in culture was previously described in Juodeikis et al.^29^ and includes the following steps: In duplicate, 20 μL of 30 μg/ml FM4-64 (Thermo Fisher Scientific) was combined with 180 μL of filtered culture supernatant or a linoleic acid standard in water (100, 75, 50, 20, 10, 5, 1, 0 μg/ml, prepared from a 1 mg/ml stock) in black 96-well plates. Following a 10-minute incubation at 37°C, endpoint fluorescence was analyzed using the FLUOStar Omega microplate reader with pre-set FM 4–64 settings (Excitation: 515-15; Dichroic: auto 616.2; Emission 720-20), employing an enhanced dynamic range. Linear standard curves from the linoleic acid samples were established for lipid quantification.

#### Proteomics

Protein profiles of whole cell lysates (WCL) and AEVs were analyzed. Therefore, 20 mg of cell biomass (3 replicates per species) were subjected to extensive ultrasonication with 400 µl of PBS. Cell debris was removed with centrifugation at 800 g at 4°C, for 5 min. The supernatants were collected for proteomic analysis. The protein content of the whole cell lysate was determined by Pierce BCA protein assay according to the manufacturer’s protocol (Thermo, USA). Protein concentration of AEVs was measured by Qubit^®^ Protein Assay (Thermo Fisher Scientific), as described above.

#### Mass spectrometry analysis

For LC-MS/MS analysis, 2 (for AEVs) or 5 µg (for WCLs) of protein were reduced and alkylated for 10 min at 95 °C with final 10 mM TCEP (tris(2-carboxyethyl)phosphine) and 40 mM CAA (2-Chloroacetamide). The sample was processed according to the SP3 protocol^36^ and digested overnight with trypsin (Promega, enzyme/protein 1:50). Peptides were desalted using SBD-RPS tips as previously described^37^. 400 ng per sample (re-dissolved in 2% acetonitrile/0.1% formic acid in water) was subjected to LC-MS/MS analysis. Protein digests were separated by nano-HPLC (Dionex Ultimate 3000, Thermo Fisher Scientific(Dionex Ultimate 3000) equipped with a C18, 5 µm, 100 Å, 100 µm x 2 cm enrichment column and an Acclaim PepMap RSLC nanocolumn (C18, 2 µm, 100 Å, 500 x 0.075 mm) (all Thermo Fisher Scientific, Vienna, Austria). Samples were concentrated on the enrichment column for 5 min at a flow rate of 15 µl/min with 0.1 % formic acid as isocratic solvent. Separation was carried out on the nanocolumn at a flow rate of 300 nl/min at 60 °C using the following gradient, where solvent A is 0.1 % formic acid in water and solvent B is acetonitrile containing 0.1 % formic acid: 0-5 min: 2 % B; 5-123 min: 2-35 % B; 123-124 min: 35-95 % B, 124-134 min: 95 % B; 134-135 min: 2 % B; 135-150 min: 2% B. The maXis II ETD mass spectrometer (Bruker Daltonics, Germany) was operated with the captive source in positive mode with the following settings: mass range: 200–2000 m/z, 2 Hz, capillary 1,600 V, dry gas flow 3 L/min with 150°C, nanoBooster 0.2 bar, precursor acquisition control top 20 (collision induced dissociation (CID). The mass spectrometry proteomics data were deposited to the ProteomeXchange Consortium^38^ via the partner repository with the dataset identifier PXD053245 (Reviewer access details: Log in to the PRIDE website using the following details: PDX accession: PXD053245;Username; Password: MFqECDz7Uyv6)^38^.

The LC-MS/MS data were analyzed by MSFragger^39,40^ by searching the public *Methanobrevibacter* protein databases (UP000232133; UP000003489; UP000004028; UP000018189; UP000001992), the archaeal protein catalogue described in Chibani et al.^22^ and a list of common contaminants^41^. Additional information on proteins found in all vesicles was retrieved via MaGe^42^ and the implemented functions SignalP (version 4.1)^43^, MHMM (version 2.0c)^44,45^ and InterProScan^46,47^, as well as from the InterPro Database^47^ (Supplementary Table 5).

Carbamidomethylation of cysteine and oxidation on methionine were set as a fixed and as a variable modification, respectively. Detailed search criteria were used as follows: trypsin, max. missed cleavage sites: 2; search mode: MS/MS ion search with decoy database search included; precursor mass tolerance ± 20 ppm; product mass tolerance ± 15 ppm; acceptance parameters for identification: 1% protein FDR^48^.

Data from EV and whole cell lysates were processed with Perseus software version 1.6.15.0. Data was filtered for decoy hits and contaminants. After log2 transformation, and subtracting the median from the column proteins were filtered for containing at least 2 valid values in at least one group.

### Mass spectrometry derived AEV metabolomics

Biological triplicates of the vesicle preparations were used for the LC-MS analysis, and a technical duplicate of a non-cultured medium that had passed through the pipeline for vesicle isolation was used as a medium blank. All samples were stored at -70°C until processing at the Vienna BioCenter Metabolomics Core Facility.

The samples were diluted with 50 μL ACN and subjected to analysis with liquid chromatography-mass spectrometry (LC-MS). 11 μL of each sample was pooled and used as a quality control (QC) sample. Samples were randomly injected on an iHILIC®-(P) Classic HPLC column (HILICON AB, 100 x 2.1 mm; 5 µm; 200 Å, Sweden) with a flow rate of 100 µL/min delivered through an Ultimate 3000 HPLC system (Thermo Fisher Scientific, Germany). The stepwise gradient has a total run time of 35 min, starts at 90 % A (ACN), and takes 21 min to 60% B (25 mM ammonium bicarbonate) followed by 5 min hold at 80% B and a subsequent equilibration phase at 90%. The LC was coupled to a high-resolution tandem MS instrument (Q-Exactive Focus, Thermo Fisher Scientific, Germany). The ionization potential was set to +3.5/-3.0 kV, the sheet gas flow to 20, and an auxiliary gas flow of 5 was used. Samples were flanked by a blank and a QC sample for background labeling and data normalization, respectively.

The obtained data set was processed by “Compound Discoverer 3.3 SP2” (Thermo Fisher Scientific). Annotation of the compounds was done through searching against our internal mass list database generated with authentic standard solutions (highest confidence level). Additionally, the mzCloud database was searched for fragment matching and ChemSpider hits were obtained using BioCyc, Human Metabolome Database, *E. coli* Metabolome Database, and KEGG databases. Only metabolites identified with highest confirmation (match with internal database) were examined in more detail; additional ones are provided in Supplementary Table 7).

The log2 fold changes, as well as p-values, were calculated by the Compound Discoverer software (Tukey HSD test (posthoc), after an analysis of variance (ANOVA) test).

### Co-incubation experiments with cell lines

#### Cytotoxicity tests of AEVs and BEVs

3-(4,5-Dimethyl-2-thiazolyl)-2,5-diphenyl-2H-tetrazolium bromide (MTT) cell viability assays were routinely performed at the end of the HT-29 cell culture assays^49^, but no significant reduction in metabolic activity could be observed for any condition used in this study (data not shown).

Additionally, CellTiter-Glo® 2.0 Cell Viability Assay (Promega) was used to investigate the cytotoxicity of AEVs on THP1-Blue cells, but no reduction in the viable cells could be detected (data not shown).

#### Confocal Microscopy

*M. smithii* ALI, M. intestini, *M. smithii* GRAZ-2, and *M. stadtmanae*-derived AEVs (1x10^11^/ml) were labeled with 5% DiO at 37°C for 30 minutes. Labeled DiO - AEVs (1x10^11^/well [10 µl]) were added to THP1-b cell monolayers cultured on collagen solution (Merck) coated 12-well chamber slides (IBIDI) overnight (16 hrs). THP1-b monocytes were previously induced to differentiate into macrophages using 150 nM PMA (Phorbol 12-myristate 13-acetate; Sigma, P8139). Samples were fixed using Pierce 4% PFA (ThermoFisher), permeabilized with 0.25% Triton X1000 (Sigma), and blocked with 10% goat serum in PBS. For nuclear visualization, cells were incubated with Hoechst 33342 (ThermoFisher), Alexa 647-Phalloidin to visualize intracellular membranes. As a second approach, AEVs were incubated with Archaea specific primary antibodies (Davids Biotechnologie GmbH, affinity purified, specific for *Methanobrevibacter* and *Methanosphaera*) and AF647 as the secondary antibody, and cells were labeled with Hoechst 33342 (ThermoFisher). Images were taken using a Zeiss LSM880 confocal microscope equipped with a 63x/1.40 oil objective. Fluorescence was recorded at 405 (blue, nucleus), 488 (green, AEVs), and 594 nm (red, intracellular membranes or AEVs). The red channel was adjusted using the ZEISS ZEN 3.9 (ZEN lite) software by the best-fit function.

#### HT-29 cytokine release

The HT-29 cytokine release assay was performed at the Institute of Molecular Biosciences, University of Graz. HT-29 (intestinal epithelial cells) were grown in T-175 tissue culture flask, containing Dulbecco’s Modified Eagle’s medium/ Nutrient F-12 (DMEM-F12) medium (Gibco, USA) supplemented with 10% fetal bovine serum (FBS), Penicillin-Streptomycin (100 μg/ml streptomycin and 100 Units/ml penicillin) and L-Glutamine (2 mM) at 37°C in a CO2 incubator. To investigate the pro-inflammatory potency of AEVs and BEVs, HT-29 cells were seeded in a 24 well tissue culture plates at a concentration of 6 x 10^5^ cells/well and cultivated for 24 h in DMEM-F12 medium supplemented with 10% fetal bovine serum (FBS), Penicillin-Streptomycin and L-Glutamine. Then, intestinal epithelial cells were washed once with PBS and the medium was replaced with AEVs or BEVs (10^8^ particles/ ml) resuspended in DMEM-F12 medium without FBS. After incubation for 20 h the cell culture supernatant was harvested, centrifuged for 2500 rpm at 4°C for 10 min to remove the cell debris and stored at -20 °C for subsequent Interleukin 8 (IL-8) quantification by ELISA, which was performed as previously described according to the manufacturer’s protocol^25^.

#### Statistics and data visualization

Vesicle properties (Concentration, size, nucleic acids, and protein content) and metabolites were plotted as boxplots in R (R-Core-Team, 2024) using the ggplot2 Package (v3.5.1)^50^.

Creation of Venn diagrams was performed by using the online tool interactiVenn^51^. PCA was created with Perseus software (v1.6.15.0)^52^.

The overview of proteins identified in archaeal vesicles and whole cell lysates, as well as proteins annotated as adhesins, were displayed in heatmaps using ggplot2 (v3.5.1) ^50^, with data transformation performed using the reshape2^53^ package (v1.4.4; Wickham, 2007). Barchart of mean intensities of protein categories was plotted with ggplot2 (v3.5.1) ^50^, and dplyr (v1.1.4)^54^ was used for the calculation of mean and standard deviation. IL-8 excretion in the HT-29 cell line was visualized as a bar chart using ggplot2 (v3.5.1) ^50^, with data transformation by reshape2^53^ (v1.4.4; Wickham, 2007), FSA (v0.9.5)^55^, and ggsignif (v0.6.4)^56^. For IL-8 excretion Kruskal-Wallis test followed by Dunn’s Multiple comparison where all EV samples were compared to the NTC (no treatment control).

## Results

Vesicles produced by Methanobrevibacter intestini*, M. smithii* ALI, *M. smithii* GRAZ-2 and *Methanosphaera stadtmanae* were visualized by electron microscopy in supernatants of cultures in the late exponential/stationary phase. Using protocols developed and optimized for BEV analysis, AEV biomass production and an isolation protocol were established to enable characterization with respect to size, composition, ultrastructure, proteome, metabolome, and interaction with mammalian cells.

### AEV formation in all methanogen species

Negative staining-and ultra-thin electron-microscopy-based methods revealed the presence of vesicle-like structures within (Fig. 1 F,I, J) and attached to the cells (Fig. 1 A, E, H, K) and in their close vicinity (Fig. 1 B) in all methanoarchaeal cultures. These were usually round shaped, approximately 87 - 198 nm in size (∼130 nm on average, sizes measured during NTA, Fig. 1 C, D, G, L), and showed a clear, sharp edge. No vesicles were observed in culture media controls that also underwent microscopy imaging.

**Figure 1:**
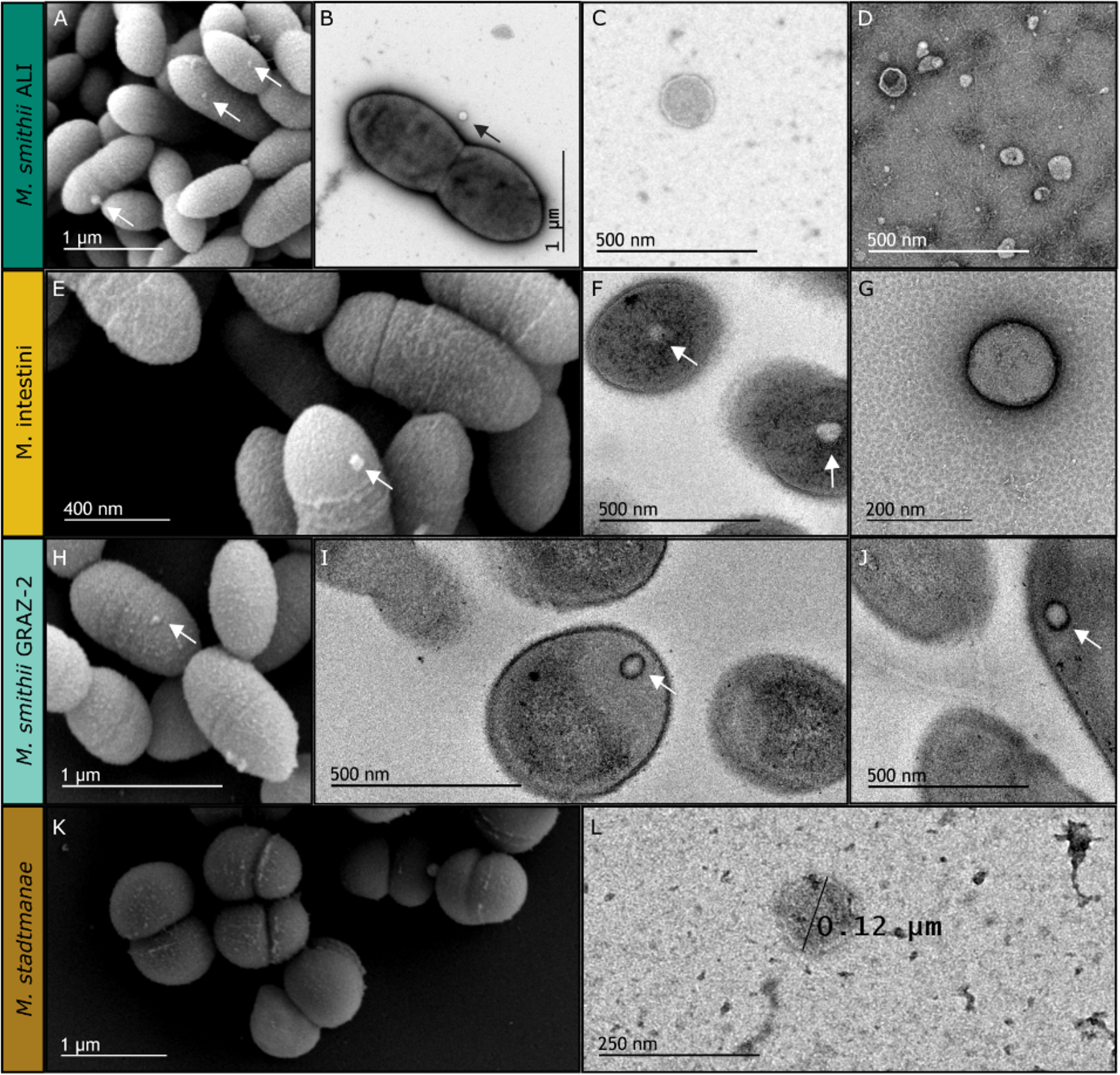
Ultrastructure of cells and vesicles. Panels A, E, H, K: Scanning electron micrographs of whole cells. B, F, I, J: Transmission electron micrographs of whole cells showing vesicles inside or attached to the cells. C, D, G, L: isolated vesicles, transmission electron micrographs. The arrows indicate the presence of AEVs.

### Biophysical AEV characteristics

AEVs from the methanogens *M. smithii* ALI, *M. smithii* GRAZ-2, M. intestini and *M. stadtmanae* were purified using a centrifugation, filtration and concentration pipeline, previously established for bacterial EVs^27^ with minor adaptations (see materials and methods). To the former described BEV isolation protocol, a centrifugation step (10.000 x g, 20 min) was added to remove residues from the culture media. The basic characteristics of the AEVs (size, concentration, nucleic acid, protein and lipid content) are summarized in Table 1 and Fig. 2. The size of the AEVs ranged from 86.9 to 197.3 nm (∼130 nm on average). M. intestini derived AEVs were found to be the largest (∼136 nm on average), and *M. smithii* GRAZ-2-derived vesicles the smallest (∼117 nm), which were similar in size to those of *M. smithii* ALI (∼124 nm). Overall the size of the vesicles was in the range of BEVs (20-400 nm)^3,57,58^ e.g. from enterotoxigenic *Escherichia coli* (ETEC, ∼120 nm), but slightly smaller than BEVs e.g. from *Bacteroides thetaiotaomicron* (∼180 nm)^59^, and *B. fragilis* (∼194 nm).

**Figure 2:**
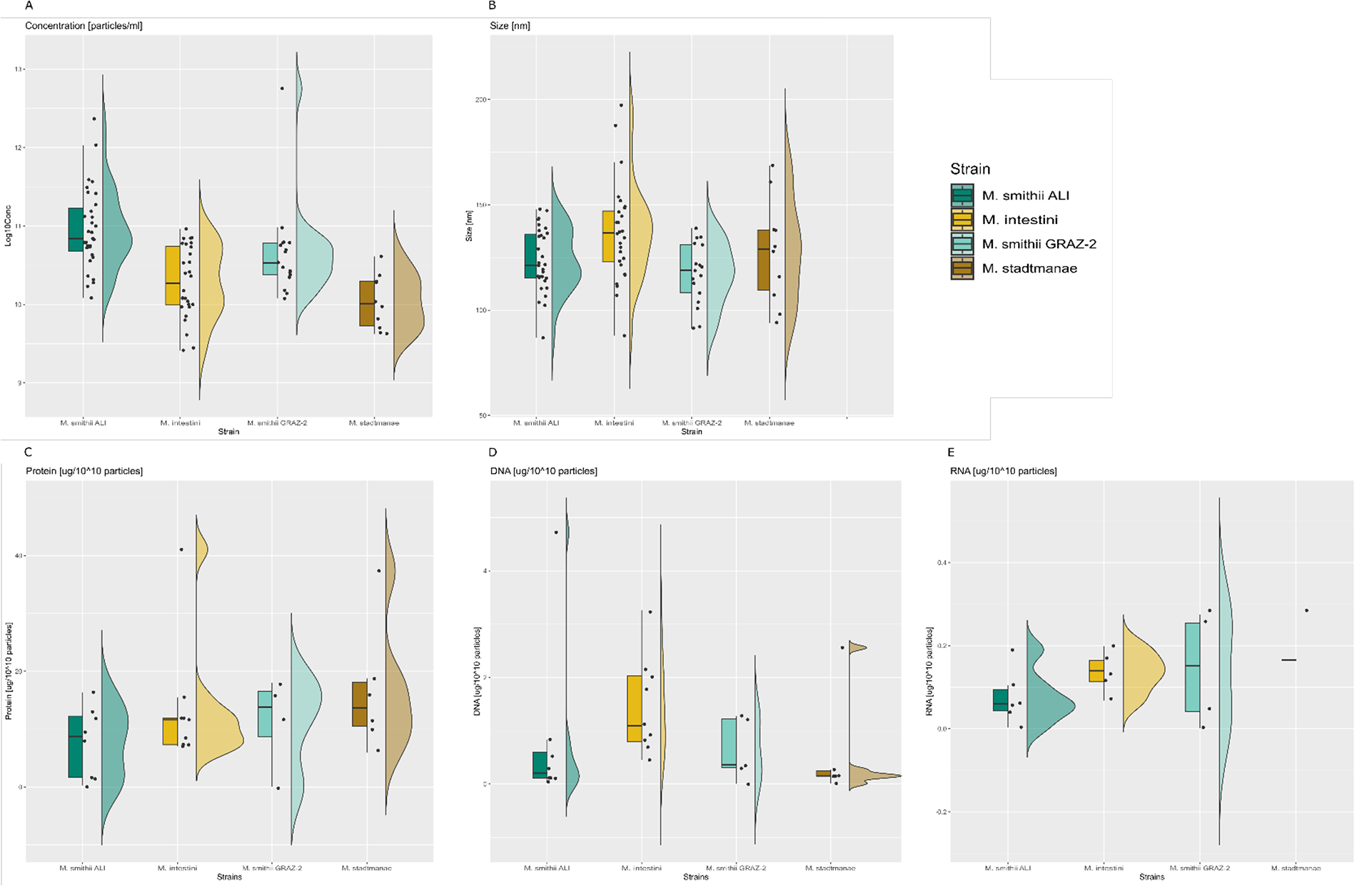
Vesicle properties of *M. smithii* ALI, M. intestini, *M. smithii* GRAZ-2, and *M. stadtmanae*. (A) Concentration [particles/ml], (B) Size [nm]. (C) Protein, (D) DNA, and (E) RNA content was normalized to [µg/10^10^ particles]. One outlier was removed in (D), and (E).

**Table 1.** Summary of archaeal vesicles’ properties, namely size, concentration, protein, DNA, RNA, and lipid content. Protein, DNA, RNA, and lipid contents were normalized to µg/10^10^ particles.

| | | Size [nm] | Concentration [particles/ml] | Protein $[\mu\text{g}/10^{10}$ particles] | DNA $[\mu\text{g}/10^{10}$ particles] | RNA $[\mu\text{g}/10^{10}$ particles] | Lipid $[\mu\text{g}/10^{10}$ particles] |
| --- | --- | --- | --- | --- | --- | --- | --- |
| *M. smithii* ALI | Mean | 123.61 | 2.16E+11 | 7.76 | 0.84 | 0.08 | 4.87 |
|  | Min. | 86.90 | 1.21E+10 | 0.26 | 0.04 | 0.04 | 1.75 |
|  | Max. | 148.00 | 2.33E+12 | 16.32 | 4.72 | 0.19 | 10.90 |
| *M. intestini* | Mean | 136.85 | 3.03E+10 | 13.89 | 3.14 | 0.14 | 81.20 |
|  | Min. | 87.90 | 2.60E+09 | 7.01 | 0.46 | 0.07 | 81.20 |
|  | Max. | 197.30 | 9.20E+10 | 41.07 | 18.27 | 0.2 | 81.20 |
| *M. smithii* GRAZ-2 | Mean | 117.28 | 3.69E+11 | 51.78 | 0.63 | 0.14 | 11.00 |
|  | Min. | 91.30 | 1.20E+10 | 0.09 | 0.004 | 0.001 | 11.00 |
|  | Max. | 138.90 | 5.63E+12 | 180.63 | 1.28 | 0.28 | 11.00 |
| *M. stadtmanae* | Mean | 127.92 | 1.44E+10 | 16.62 | 0.55 | 0.17 | 35.30 |
|  | Min. | 94 | 4.20E+09 | 5.93 | 0.01 | 0.17 | 35.30 |
|  | Max. | 168.70 | 4.08E+10 | 37.33 | 2.56 | 0.17 | 35.30 |

Average concentration of the retrieved AEVs (Table 1) was much lower than concentrations usually measured for BEVs, such as ETEC (6.38E+11 particles/ml), and *B. fragilis* (8E+11 particles/ml). The concentrations were reasonably consistent for all *M. smithii* strains, with lower concentration retrieved for M. intestini and *M. stadtmanae* (Table 1).

Overall, the protein content ranged from 0.09 to 180.6 µg/10^10^ particles. On average, AEV extracts of *M. smithii* GRAZ-2 contained the highest protein concentration (∼ 52 µg/10^10^ particles), whereas the lowest concentrations were found in *M. smithii* ALI (∼7.8 µg/10^10^ particles). Regarding the lipid content, all AEVs extracts were in the range of the standard linoleic acid (20-100 µg/ml). AEVs from M. intestini showed the highest (∼81.20 µg/10^10^ particles) and the lowest amount of lipids (∼4.9 µg/10^10^ particles on average). It has to be mentioned that the lipid content could only be detected in a few samples, as not all measurements of concentrations were found in the standard range. Overall, the DNA content of AEV extracts ranged from 0.004 to 18.27 µg/10^10^ particles. AEVs of M. intestini had the highest DNA concentration (3.14 µg/10^10^ particles), while *M. stadtmanae* had the lowest one (0.55 µg/10^10^ particles) on average. RNA content of AEVs could not be detected in all samples, due to low concentrations. AEVs from *Methanobrevibacter* strains contain similarly low amounts of RNA (0.08 - 0.014µg/10^10^ particles), while 0.17 µg/10^10^ particles could be detected for *M. stadtmanae*.

### Methanobrevibacter AEVs have comparable proteomes and show a massive enrichment in adhesins

The protein cargo of *M. smithii* ALI and M. intestini EVs were compared with that of their respective whole microbial cell proteomes (whole cell lysate, WCL). Profiling was carried out through LC-MS/MS, employing isolated AEVs and whole cell lysates (n=3) of *M. smithii* ALI and M. intestini (refer to material and methods section for details). A total of 1475 vesicular proteins across all isolated EVs were identified (*M. smithii* ALI: 801; M. intestini: 674), complemented by the identification of 2537 proteins from the whole cell lysates (WCL; *M. smithii* ALI: 1262; M. intestini: 1275, Supplementary Fig. S3, Supplementary Table 1). Proteins were considered to be present in a sample, based on a prevalence in three out of three replicates for each group of sample (WCL *M. smithii* ALI (WCL_ALI): 1026; WCL M. intestini (WCL_int): 1100; EVs M. smithii ALI (EV_ALI): 364, and EVs M. intestini (EV_int): 259; Supplementary Fig. S2, Supplementary Table 1). Compared to the 2047 proteins identified in BEVs derived from *B. thetaiotaomicron*^59^, the total number of proteins in methanoarchaeal EVs was much lower.

AEVs derived from *M. smithii* ALI (EV_ALI) and M. intestini (EV_int) share 229 proteins, while having 135 and 30 unique proteins in these strains, respectively (Fig. 3 B). Only a small number of proteins (EV_ALI: 35, EV_int: 56) were detected in the vesicles but not in the whole cell lysates (Fig 3 D and E). Proteins of whole cell lysates were highly similar, as 816 proteins were identified in both WCL_ALI and WCL_int (Fig 3 A). 173 proteins were found in all four groups (EV and WCL of both strains, Fig 3 C).

**Figure 3:**
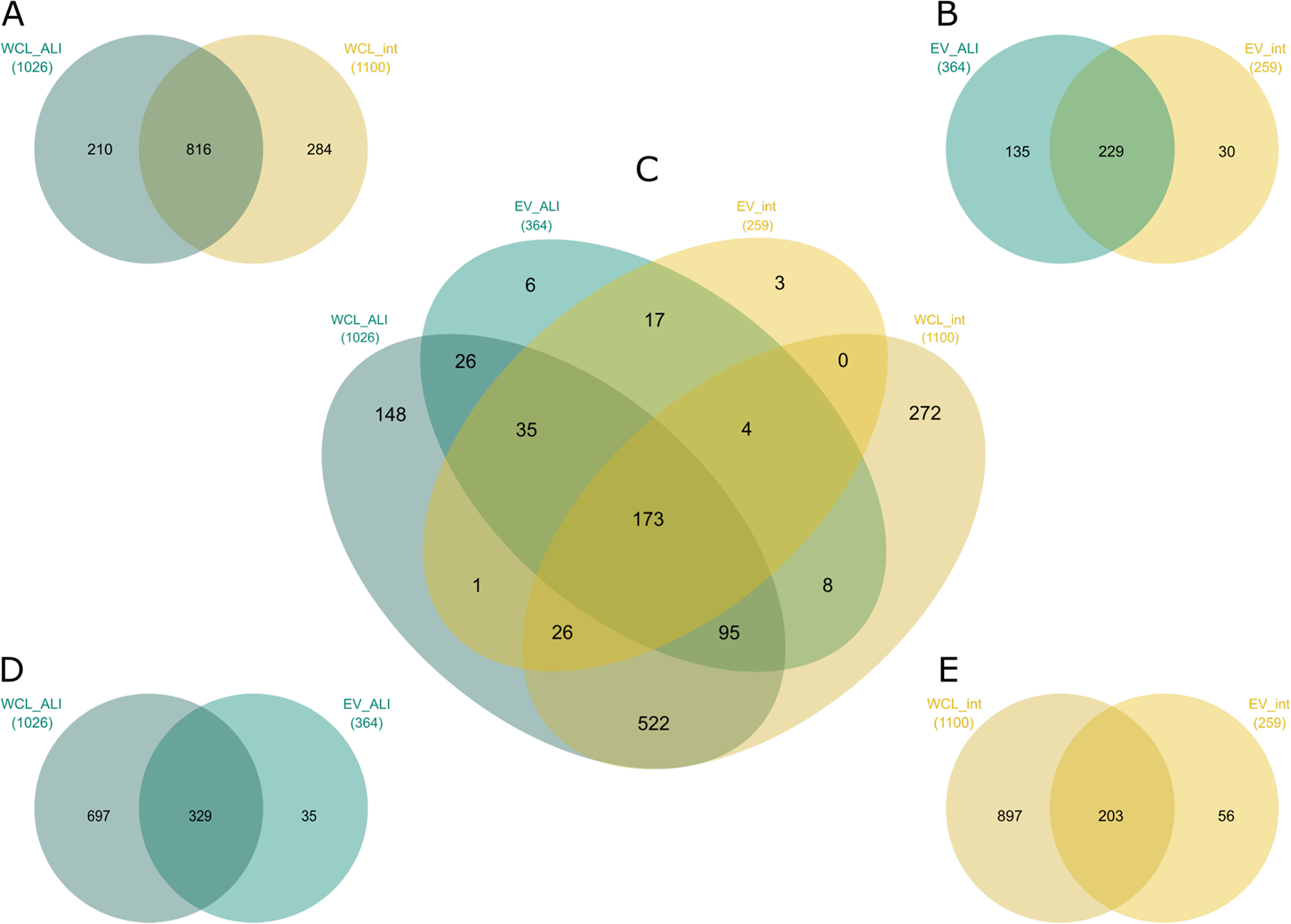
Venn diagrams constructed with proteins based on a prevalence in 3 of 3 replicates per group of WCL_ALI, WCL_int, EV_ALI or EV_int. (A) WCL_ALI vs. WCL_int, (B) EV_ALI vs. EV_int, (C) WCL_ALI vs. EV_ALI vs. EV_int vs. WCL_int, (D) WCL_ALI vs. EV_ALI, and (E) WCL_int vs. EV_int. EV_ALI, vesicles M. smithii ALI; EV_int, vesicles M. intestini; WCL_ALI, whole cell lysate *M. smithii* ALI; WCL_int, whole cell lysate M. intestini.

The PCA plot depicted in Figure 4 A illustrates different distribution patterns between whole-cell lysates (WCL) and extracellular vesicles (EVs) for both strains. Notably, it also highlights the similarities observed between WCL_ALI and WCL_int, as well as between EV_ALI and EV_int.

**Figure 4:**
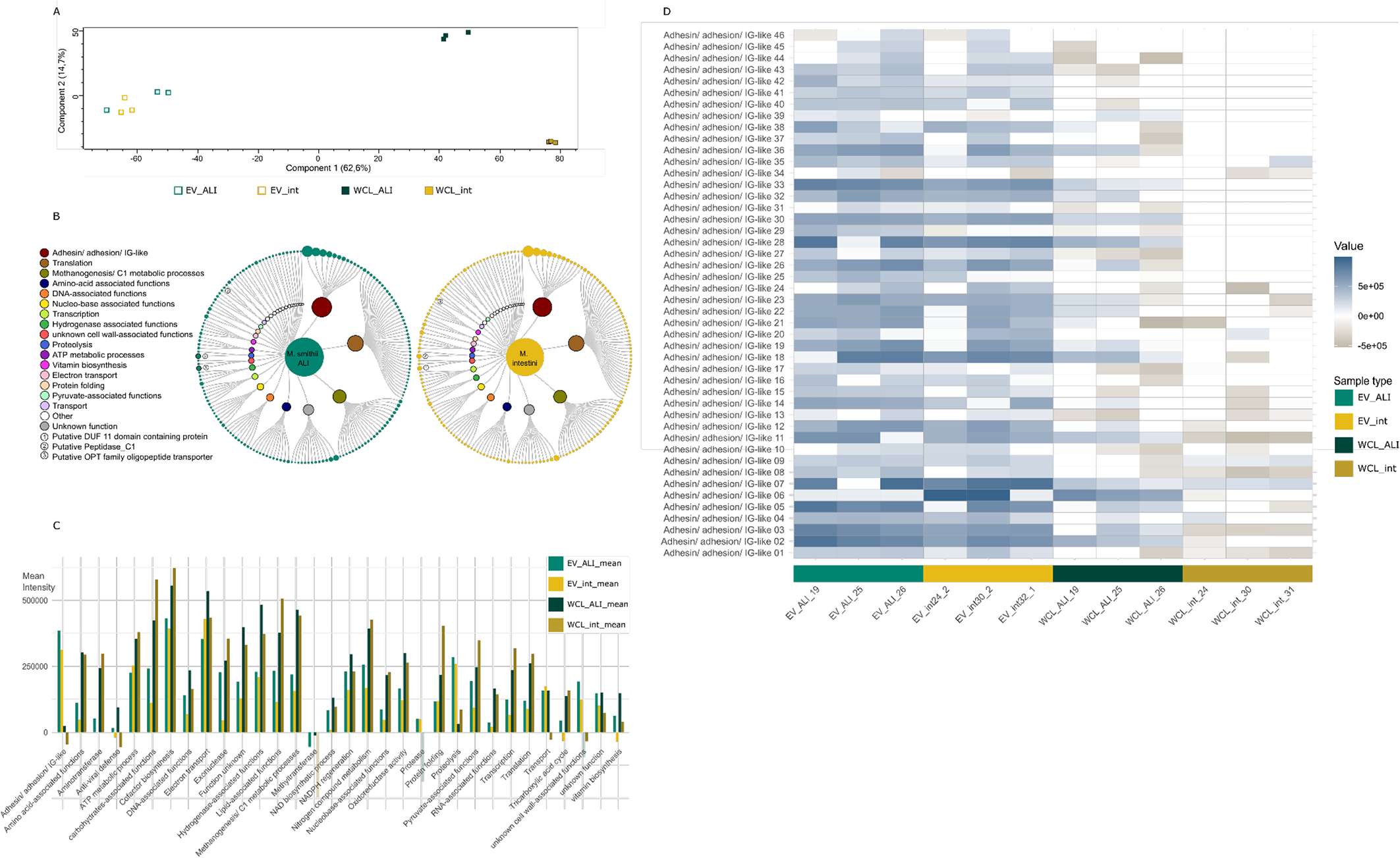
(A) PCA plot, proteins prevalent in 3 of 3 replicates per group. EV_ALI, vesicles *M. smithii* ALI; EV_int, vesicles M. intestini; WCL_ALI, whole cell lysate *M. smithii* ALI; WCL_int, whole cell lysate M. intestini. (B) 229 overlapping proteins found in the proteomes of *M. smithii* ALI (left, n=3 biological replicates) and M. intestini (right, n=3 biological replicates) vesicles, grouped by their intensities (reflected by the size of the circles) and (putative) functions (details can be retrieved from Supplementary Table CME-2). Visualization was done via RawGraphs^60^ and InkScape^61^ (C) Barchart showing the mean intensities of protein categories in vesicles (EV) and whole cell lysates (WCL). Only proteins which were found in 3+3 biological replicates of *M. smithii* ALI (ALI) and M. intestini (int) (n=229) are included (Data: Supplementary Table 4). (D) Heatmap depicts the presence of 46 proteins annotated as adhesin/adhesion/IG-like present in six out of six AEV extracts (EV_ALI, EV_int) compared to the whole cell lysate (WCL_ALI, WCL_int). EV_ALI, vesicles *M. smithii* ALI; EV_int, vesicles M. intestini; WCL_ALI, whole cell lysate *M. smithii* ALI; WCL_int, whole cell lysate M. intestini.

The protein content of the vesicles of both *Methanobrevibacter* species was strikingly similar (Fig. 4, Supplementary Fig. S2 and S3), with 229 proteins found in all six extracts. The most abundant proteins were Adhesins/ adhesin-like proteins/ proteins with an IG-like domain, as identified through InterPro prediction (ALPs, Fig. 4 B and C; for details on functional annotation see Materials and Methods and Supplementary Table 6); these proteins were also highly enriched compared to the whole-cell lysates (WCL, Fig. 4 B and C; Supplementary Table 4)^47^. ALPs are rarely studied in archaea, but were found to be very abundant in e.g. rumen methanogens where they account for up to 5% of all genes. It has been suggested that the ALPs serve to attach to their protozoan hosts or to the cell surface of bacteria^62^. ALPs have also been found in human-associated *Methanobrevibacter* species^63^, for which adhesion and sugar-binding function has been proposed. Indeed, the identified vesicle-associated ALPs carried a variety of protein domains, indicative of an adhesive (Invasin/intimin cell-adhesion fragments; IG-like_fold superfamily) and polysaccharide binding functions (PbH1; pectin_lyase_fold, Pectin_lyase_fold/virulence; details for all genes and their identified domains are given in Supplementary Table 5).

Bacterial proteins containing IG-like domains exhibit a broad spectrum of functions, such as cell host adhesion and invasion. IG-like domains are also found in periplasmic chaperones and proteins that assemble fimbriae, in oxidoreductases and hydrolytic enzymes, ATP-binding cassette transporters, sugar-binding and metal-resistant proteins^64^. These proteins are structural components of bacterial pilus and nonpilus fimbrial systems and members of the intimin/invasin family of outer membrane adhesins, indicating their relevance for adhesion and interaction with the biological surroundings. Microbial pectin and pectate lyases are involved in the degradation of pectic components of the plant cell, which is an important trait for plant pathogens, as well as the degradation of dietary components in the gastrointestinal tract. However, this specific β-helix topology has various functions e.g. as galacturonases, or for the adhesion to mammalian cells^65^.

Within a group of transport-associated proteins, we found substantial enrichment of a protein (representative: GUT_GENOME043902_01504) with an OPT (oligopeptide transporter) superfamily domain, which in prokaryotes may contribute to iron-siderophore uptake^66^, indicating a potential role in iron binding.

A further substantial increase was observed for a putative DUF11 domain-containing protein^67^, which might be important for stabilizing surface wall structures in *Methanothermobacter* sp. strain CaT2 ^67^. Another interesting finding was the increased presence of a putative peptidase_C1, which also showed adhesin-like domains (Supplementary Table 4 and 6).

### Metabolite cargo of AEVs could have an effect on gut-brain-axis

Similar to the proteomic analyses, the metabolic profiles of AEVs of *M. smithii* ALI and M. intestini were overall similar, but with high variability across biological replicates, probably due to variations in input concentrations (see group CV % in Supplementary Table 7; Fig. 4).

Strikingly, the AEVs of M. intestini revealed a significantly increased content of glutamic and aspartic acid (Fig. 5; *P* =0.03 and *P* =0.01, respectively; Supplementary Table 7; this table also includes details on statistics). Also, the AEVs of *M. smithii* ALI revealed a substantial Log2 Fold Change (FC) compared to background samples, indicating that these amino acids are important cargos for both species. *M. smithii* ALI were substantially loaded with arginine.

**Figure 5:**
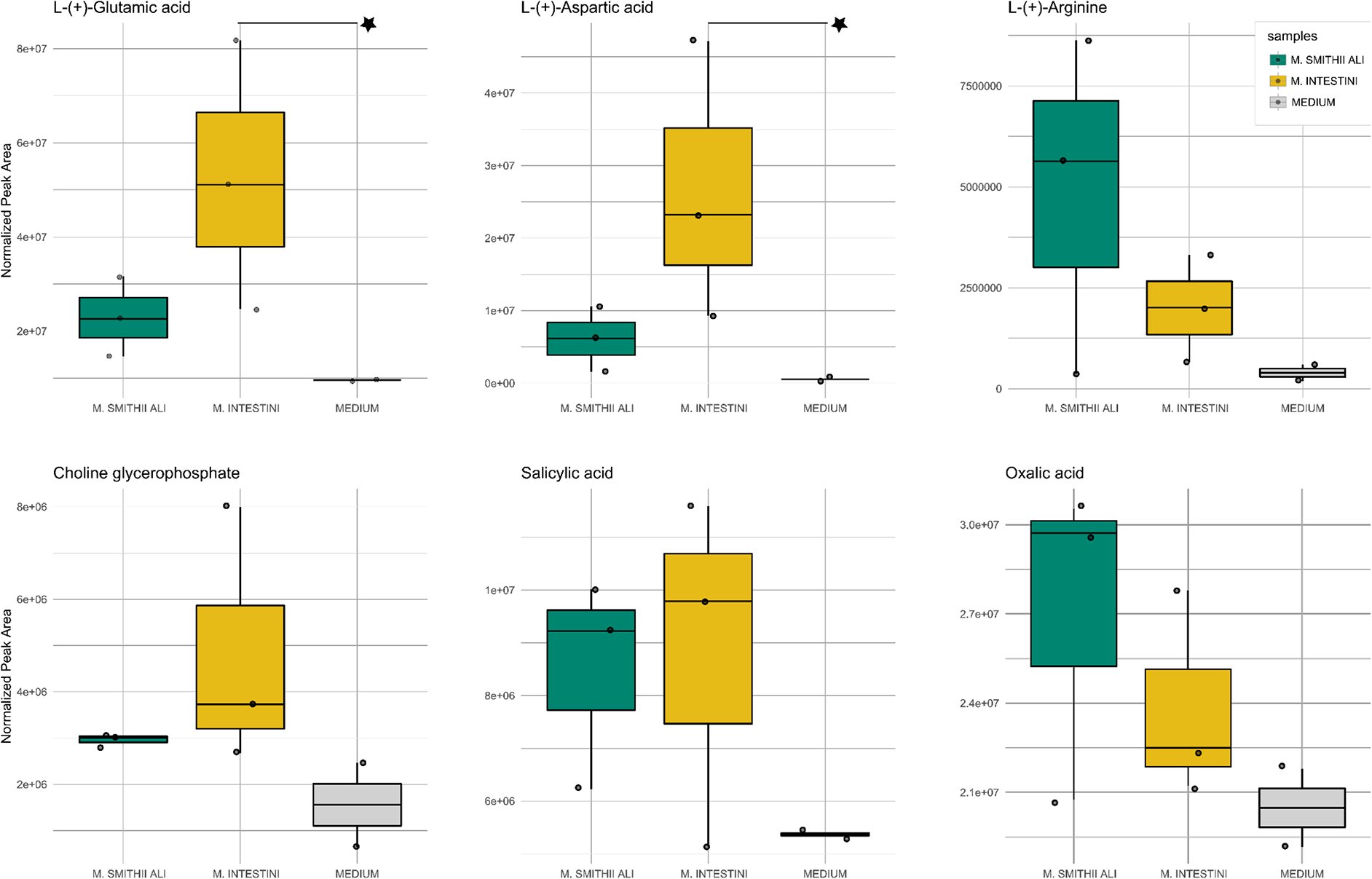
Metabolomics: Metabolites detected in archaeal vesicles (biological triplicates), compared to the culture medium control (technical duplicates). The Y-axis shows the normalized peak area (LC-MS). Significantly changed compounds are highlighted by an asterisk.

Notably, glutamate has been identified as a component of bacterial EVs (*B. fragilis)*^68^. Besides their roles in central metabolism, both amino acids are considered to act as neurotransmitters^69^. Glutamate plays a fundamental role as an excitatory neurotransmitter in the central nervous and the enteric nervous system and acts, together with other metabolites, along the “microbiota-gut-brain axis”^70^ as an “interkingdom communication system”. It is considered that the glutamatergic receptors, along the microbiota-gut-brain axis, could have an impact on multiple physiological responses in the brain and gut. As glutamate usually does not enter the bloodstream from the large intestine, AEVs could be supporting the transmission to glutamatergic enteric neurons/ receptors^70^. Despite its potential function as a neurotransmitter, aspartate also supports the proliferation of mammalian cells (e.g. cancer cells)^59^.

Choline glycerophosphate (glycerophosphorylcholine, alpha GPC) was found to be elevated in AEVs of both species (Figure 5). Also, for this compound, a potential neurological effect was described, which has been considered for the treatment of Alzheimer’s disease^71^.

The origin of the salicylic acid, which was found to be increased in AEVs of both species, is unclear (potentially from chorismate), but its potential effects on the host and microbiome could include bactericidal and antiseptic action in higher concentrations^72^. Another compound found to be increased was oxalic acid, the latter having the characteristics of a chelating agent for metal cations, making insoluble iron compounds into a soluble complex ion, which could be an interesting trait for gastrointestinal microbiota^73^.

### Human monocytes acquire AEVs

Human leukemia monocytic THP-1 cells are a common model for studying monocyte/macrophage functions, signaling pathways, mechanisms, and drug and nutrient transport^74^. For visualizing the association or interaction of AEVs with host cells, AEVs of *M. smithii* ALI, *M. smithii* GRAZ-2, M. intestini, and *M. stadtmanae* were incubated with macrophage monolayers (THP1-b) for 24 h and their localization was assessed by immunofluorescence microscopy. Co-localization of DiO-labeled AEVs (Fig. 6 A-E, green dye) with and within host cell nuclei were investigated using the nuclei marker Hoechst 33342 (blue), and the cytoskeleton marker Alexa 647-Phalloidin (red). AEVs from all strains were shown to be in close association with the nuclei. Similar localization of EVs was previously described for bacterial EVs e.g. from *B. thetaiotaomicron*^75^. A representative z-stack of *M. stadtmanae* AEVs and macrophage monolayers supports the uptake of AEVs by the macrophage cells (Fig. 6 E).

**Figure 6:**
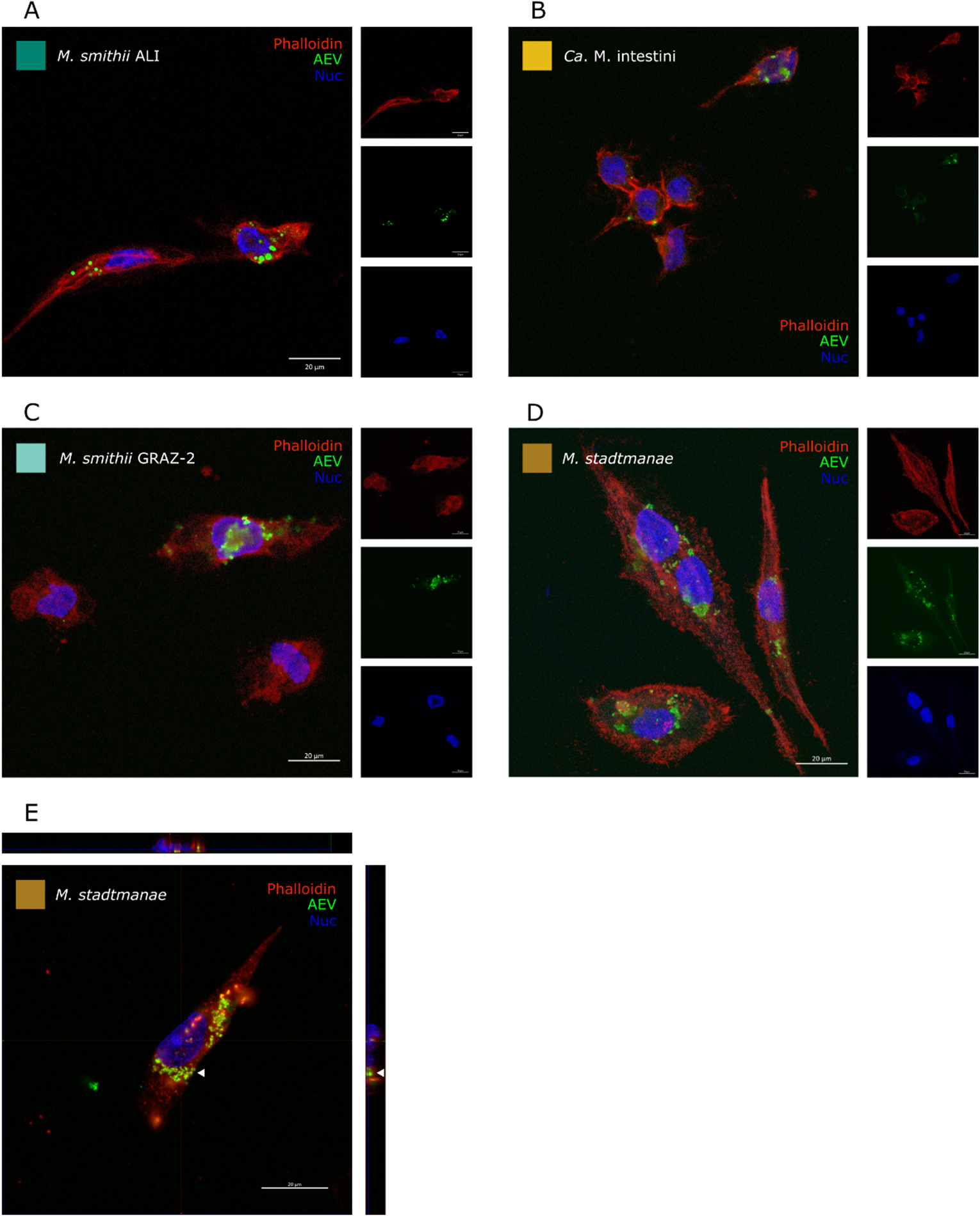
Immunofluorescence microscopy of DiO-labeled AEVs (green) acquired by human macrophages (24h incubation). Macrophage monolayers were stained with antibodies to visualize cytoskeleton (Alexa 647-Phalloidin, red), nuclei (Hoechst 33342, blue). Macrophage incubation with (A) *M. smithii* ALI derived EVs, (B) M. intestini derived EVs, (C) *M. smithii* GRAZ-2 derived AEVs, (D) *M. stadtmanae* derived AEVs. (E) Representative z-stack of *M. stadtmanae* deriver AEVs acquired by a macrophage.

### AEVs of M. intestini induce substantial IL-8 excretion in HT29 epithelial cell line

Human intestinal HT-29 cells are useful for epithelial cell research, and have recently been used in a comparative study to assess the differential pro-inflammatory potency of BEVs derived from gut bacteria^25^. To investigate the pro-inflammatory potential of AEVs, we examined the IL-8 cytokine response in intestinal epithelial cells upon exposure to AEVs derived from *M. smithii* ALI, *M. smithii* GRAZ-2, M. intestini, and *M. stadtmanae*. The pro-inflammatory cytokine IL-8 was chosen as it demonstrated a robust BEV-dependent induction in a previous study ^25^. Moreover, BEVs of enterotoxigenic *Escherichia coli* (ETEC) and *B. fragilis* were included as representatives of intestinal BEVs known to induce a very high or no IL-8 response ^25^. In concordance to a recent report^76^, AEVs derived from *M. smithii* ALI, *M. smithii* GRAZ-2, and *M. stadtmanae* failed to induce a significant increase of the IL-8 levels compared to the no treatment control (NTC, *P* > 0.05, Fig. 7). In contrast, exposure of HT-29 cells to AEVs derived from M. intestini resulted in a significant IL-8 induction (*P* < 0.001) at similar levels as observed for BEVs from ETEC (*P* < 0.001). These results suggest that AEVs derived from different archaeal species demonstrate differential pro-inflammatory potency in HT-29 cells with AEVs from M. intestini inducing a relatively strong IL-8 response.

**Figure 7:**
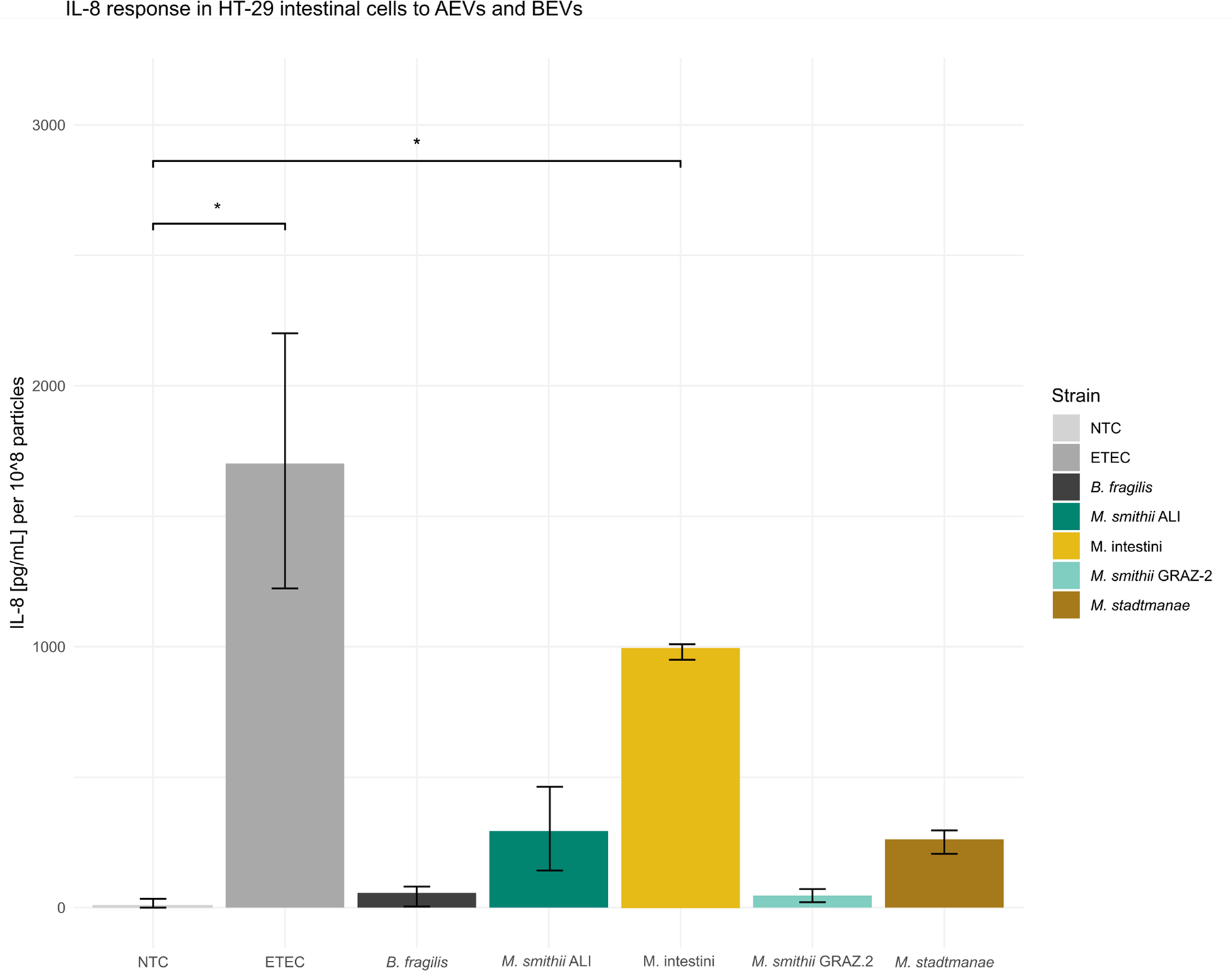
IL-8 response in HT-29 intestinal cells to AEVs and BEVs. Cytokine levels were quantified by ELISA in supernatants of HT-29 intestinal cells incubated for 16 h with equal particle amounts of AEVs or BEVs. Donor strains of the AEVs or BEVs are indicated on the X-axis. Incubation with PBS served as no treatment control (NTC). Data are indicated as the median ± interquartile range (*n* = 6). Asterisks highlight significant differences to the NTC (*, *P* < 0.001 by Kruskal-Wallis test, followed by Dunn’s *post hoc* test).

## Discussion

The discovery of archaeal extracellular vesicles (AEVs) produced by human GIT-associated archaea introduces a novel principle in archaea-microbiota and archaea-host interactions. Just like bacterial extracellular vesicles (BEVs) known from the human gut microbiota, AEVs are membrane-bound structures that transport various biomolecules, including proteins, lipids, and nucleic acids^4,8–12^. We assume that these vesicles as well, have the potential to modulate the microbial community and host physiology, by acting as a communication and cargo vehicle.

This study demonstrates that AEVs derived from GIT-associated archaea are comparable in size to BEVs, although the particle count was substantially lower for archaea (Fig. 2, Table 1). Previous research has indicated that growth conditions, such as growth stage and medium composition, can influence the particle count, size, and vesicle cargo of BEVs leading to a heterogeneity among BEVs^13,29,77–82^. It is likely that similar effects occur with AEVs. As we used biological replicates in our experiments, a certain fluctuation in e.g. metabolite cargo was observed. Heterogeneity implies that different vesicle subtypes may carry distinct cargo, potentially leading to varied biological effects by targeting different host cells or microbial cells^83–85^. Moreover, the isolation process itself might impact the retrieval of different vesicle subtypes^83^.

Both studied archaeal genera (*Methanobrevibacter* and *Methanosphaera*) were capable of vesicle formation. All vesicles were found to be acquired by human monocytes (Fig. 6), and stimulate IL-8 secretion (Fig. 7, P > 0.05). For detailed proteomic and metabolomic studies, this study focused on vesicles from *Methanobrevibacter* species, the most abundant archaea in the human gut microbiome, comprising up to 4% of the microbiome^86^. *Methanobrevibacter* species rely on syntrophic bacterial partners that provide small organic compounds like H_2_ (or formate) and CO_2_ for methanogenesis^63,87,88^. The bacterial partner benefits from this interaction, as potentially inhibiting end products of fermentation are efficiently removed^63,87,88^. As such, a well-regulated and controlled interaction with bacterial syntrophic partners is highly crucial for *Methanobrevibacter* species.

Adhesins, which were found to be highly accumulated in archaeal vesicles, have been identified to be important communication vehicles. For instance, *Methanobrevibacter* influences the metabolism of *Christensenella minuta*, shifting short-chain fatty acid (SCFA) production from butyrate to acetate^89^. This complex communication system, regulating the metabolic processes of both partners, is believed to be mediated by *Methanobrevibacter* surface adhesins, leading to significant physiological changes in the involved microorganisms^89^. From the bacterial kingdom, numerous adhesins are known to mediate interaction, colonization, infection and host interaction, making them key targets in bacterial pathogenesis^90,91^. Considering that adhesins are highly enriched in AEVs, as shown Figure 4 B-D, the importance of AEVs for archaeal-bacterial and archaeal-host interactions over longer distances becomes evident.

In *Methanobrevibacter ruminantium*, a prevalent *Methanobrevibacter* species in ruminants, 5% of the genome encodes adhesins. Among them, adhesin Mru_1499 has been identified as a crucial factor allowing *M. ruminantium* M1 to bind and interact with hydrogen-producing protozoa and bacteria (i.e. *Butyrivibrio proteoclasticus*) in the rumen, facilitating efficient methane production^62^. Other adhesins facilitate adhesion to host cells and tissues, allowing microorganisms to establish and persist within the host environment.

Next to the upregulation of adhesins upon syntrophic interactions with hydrogen-producing microorganisms, adhesins were found to be increased also under nicotinic acid limitation (vitamin B3)^62,92,93^, indicating a complex interplay of metabolite-availability and the need for interaction with the microbial community and/ or the host.

Enriching adhesins on mobile vehicles such as AEVs offers numerous benefits, including the ability to reach communication partners beyond the immediate physical proximity of the non-motile archaeal cells potentially enabling even a global regulation of bacterial metabolism.

It must be considered that also the host is a target of the AEVs, as indicated by the efficient uptake of AEVs in human monocytes, and the profound response of epithelial cells (Fig. 6 and 7). It shall be mentioned that archaeal adhesins are believed to be heavily glycosylated^94^. Glycosylation is often species-specific, which could explain the different responses of HT-29 cells to AEVs from *Methanobrevibacter smithii* ALI and Methanobrevibacter intestini, despite similar overall AEV assembly (Fig. 2, 3, and 4). This highlights the importance of studying adhesin glycosylation patterns to understand their role in host-microbe interactions.

The metabolic profiling of AEVs indicates increased levels of aspartic acid and glutamate (Fig. 5), which is intriguing and warrants further investigation. These findings suggest a potential link between archaeal AEVs and the gut-brain axis (as discussed in the results section), opening new avenues for research into how these vesicles might influence host physiology and neurological processes.

In summary, the identification of AEVs and their components provides significant insights into the complex interactions within the gut microbiome, highlighting the critical role of *Methanobrevibacter* adhesins in microbial communication and host interaction. This understanding could pave the way for novel therapeutic strategies targeting microbial interactions and their impacts on host health.

## Conclusion

Recent investigations have expanded our understanding of EVs beyond the bacterial domain, revealing their presence and significance in other microbial realms. Notably, the human archaeome, comprising archaeal communities inhabiting various niches within the human body, has emerged as a newfound player in the EV landscape. Archaea, once predominantly studied in extreme environments, have now been recognized as integral components of the human microbiota, exerting subtle yet profound influences on human health and disease.

The revelation of EV production by the human archaeome introduces a new dimension to our comprehension of microbial communication within the human body. While the specific roles and functions of archaeal EVs remain largely unexplored, their existence suggests an intricate network of interdomain interactions shaping the dynamics of the human microbiome. Furthermore, the similarities and distinctions between bacterial and archaeal EVs present intriguing avenues for comparative studies, offering insights into the evolutionary origins and adaptive strategies of extracellular vesicle-mediated communication in diverse microbial taxa.

## Acknowledgements

We thank Stefanie Duller for providing electron micrographs. The support for V. Weinberger through the local dissertation program MolMed is acknowledged. We acknowledge the JIC Bioimaging facility and staff for their contribution to this publication.

## Funding

This research was funded in whole or in part by the Austrian Science Fund (FWF) [10.55776/F83, 10.55776/P32697, and 10.55776/COE7]. The authors acknowledge the support of the ZMF Galaxy Team: Core Facility Computational Bioanalytics, Medical University of Graz, funded by the Austrian Federal Ministry of Education, Science and Research, Hochschulraum-Strukturmittel 2016 grant as part of BioTechMed Graz. The Vienna BioCenter Core Facilities (VBCF) Metabolomics Facility acknowledges funding from the Austrian Federal Ministry of Education, Science & Research; and the City of Vienna.

## Author contributions

The study was designed by CME and VW. VW, PM, and TZ isolated the vesicles, together with help from RS, EJ, and SRC. Vesicle biophysical characterization was done by VW. VW and BD performed proteomics and analyzed the data with the supervision of HK and CME. Metabolomics was performed by TKoe and GG, and data were analyzed by VW and CME. Electron microscopy was performed by DP, KH, DK, and KG. HT and SS performed experiments with HT-29 cells. Experiments with macrophages were performed by VW, with the help of RS and EJ. The lipid assay was performed by RJ. VW and CME wrote the manuscript, and TS, CK, RM, TKue, and TW contributed to the writing of the manuscript and figure preparation. The manuscript was read and approved by all authors.

## Competing interests

None declared.

## Materials & Correspondence

Material requests and correspondence should be sent to

## Notes

### Competing Interest Statement

The authors have declared no competing interest.

