## Supplementary Figures for "Proteomic and Metabolomic Profiling of Archaeal Extracellular Vesicles from the Human Gut"

Supplementary Information

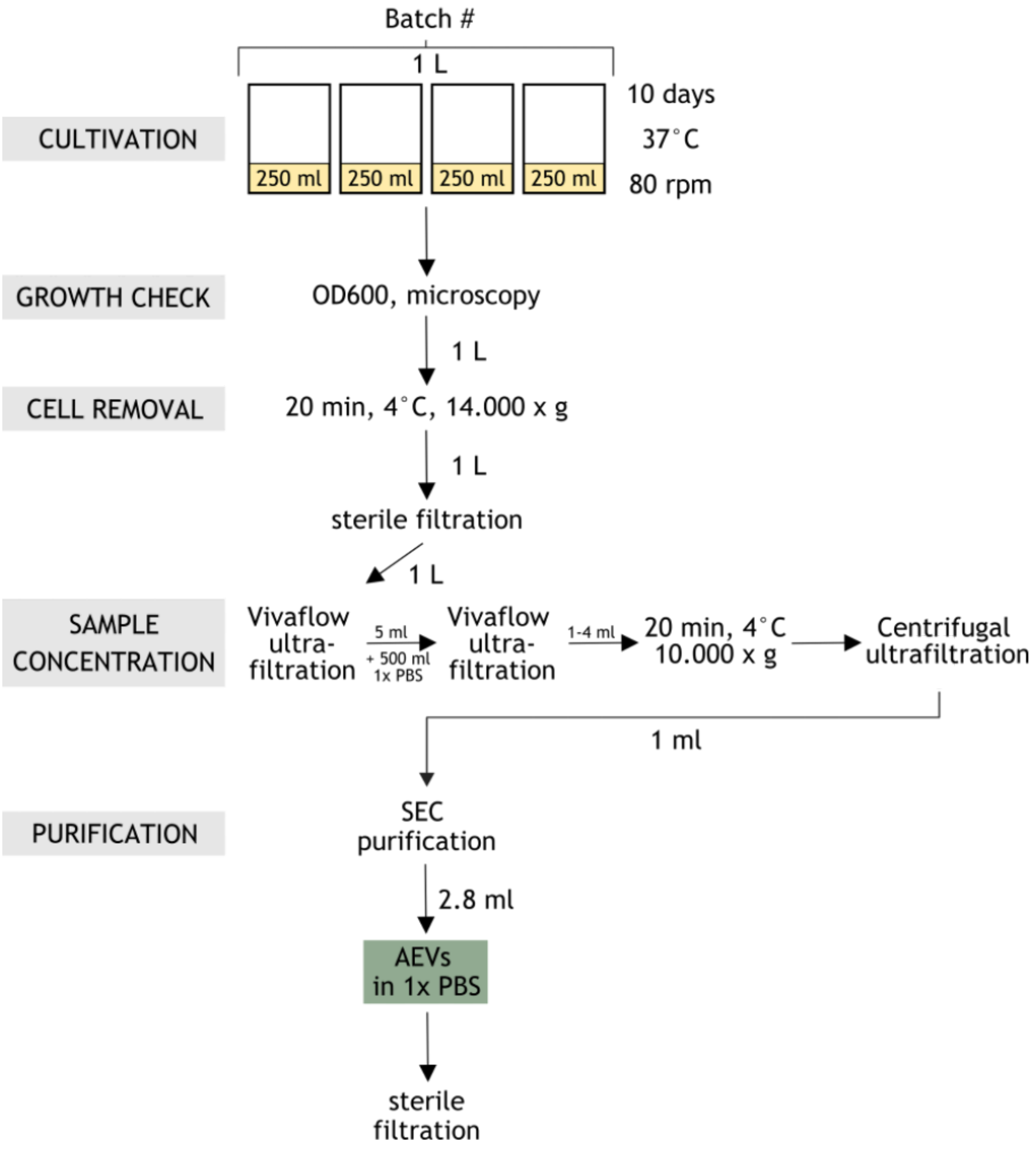


Supplementary Figure S1: Workflow of archaeal extracellular vesicle (AEV) preparation and isolation.

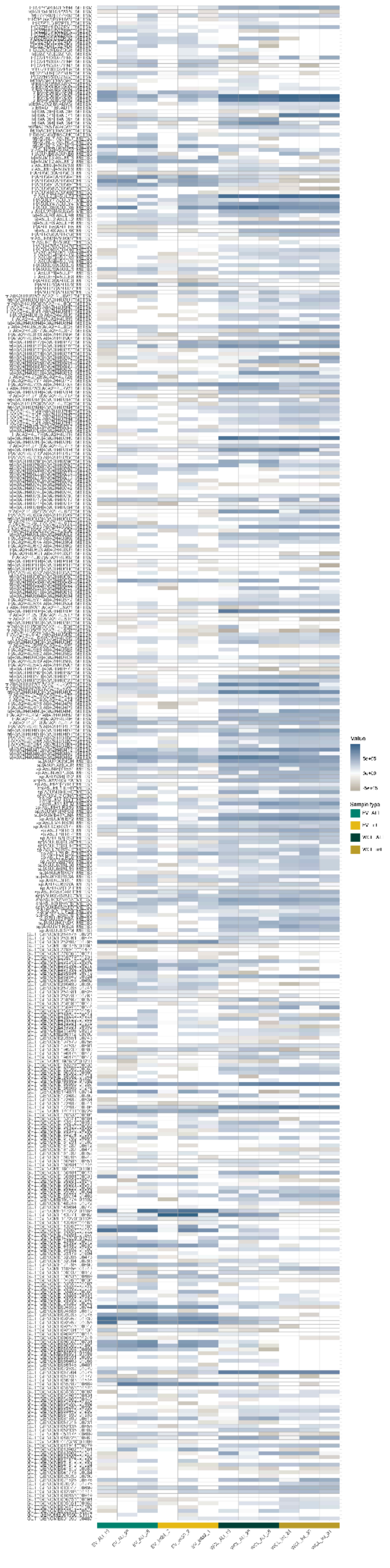

Supplementary Figure S2: Heatmap of 394 proteins based on a prevalence in 3 of 3 replicates for EV_ALI and/or EV_int. EV_ALI, vesicles *M. smithii* ALI; EV_int, vesicles M. intestini; WCL_ALI, whole cell lysate *M. smithii* ALI; WCL_int, whole cell lysate M. intestini.

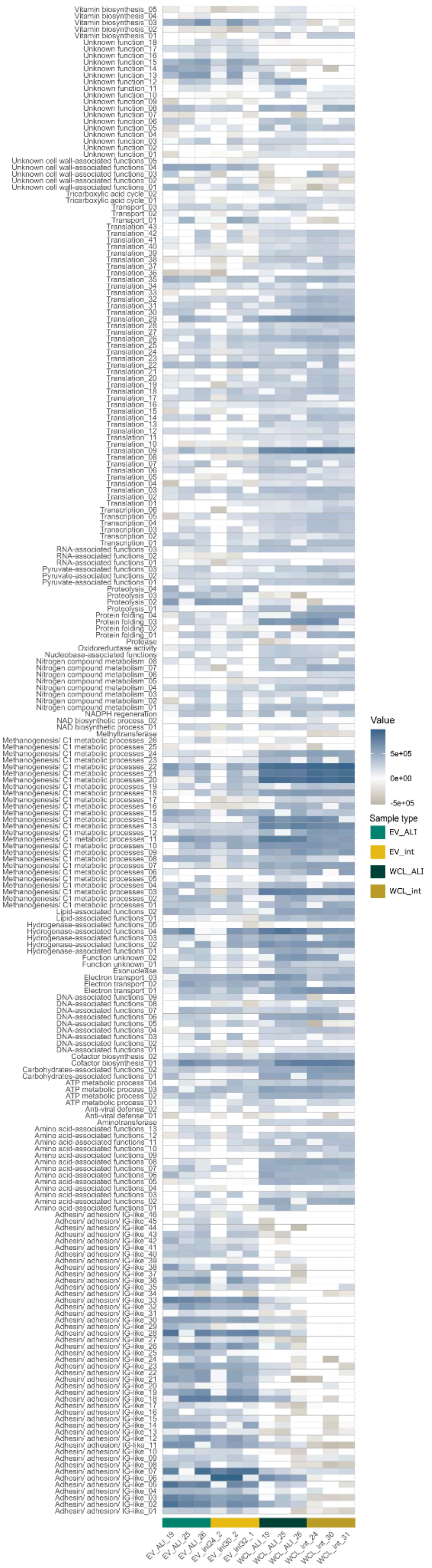


Supplementary Figure S3: Heatmap summarizes 229 proteins identified in 6 out of 6 replicates of EV samples, EV_ALI and EV_int, with their annotations. EV_ALI, vesicles *M. smithii* ALI; EV_int, vesicles M. intestini; WCL_ALI, whole cell lysate *M. smithii* ALI; WCL_int, whole cell lysate M. intestini.
